# Temporal Tracking of Plasma Cells *in vivo* Using J-chain CreERT2 Reporter System

**DOI:** 10.1101/2023.12.02.569736

**Authors:** Timothy C. Borbet, Kimberly Zaldaña, Anastasia-Maria Zavitsanou, Marcus J. Hines, Sofia Bajwa, Tate Morrison, Thomas Boehringer, Victoria M. Hallisey, Ken Cadwell, Sergei B. Koralov

## Abstract

Plasma cells (PCs) are essential for humoral immunity, as they are responsible for the production of antibodies and contribute to immunological memory. Despite their importance, differentiating between long-lived and short-lived PCs *in vivo* remains a challenge due to a lack of specific markers to distinguish these populations. Addressing this gap, our study introduces a novel J-chain CreERT2 GFP allele (IgJ^CreERT2^) for precise genetic studies of PCs. This model takes advantage of PC-restricted expression of the J-chain gene, enabling temporal and cell-specific tracking of PCs utilizing a tamoxifen-inducible Cre recombinase. Our *in vitro* and *in vivo* validation studies of the inducible Cre allele confirmed the fidelity and utility of this model and demonstrated the model’s ability to trace the long-lived PC population *in vivo* following immunization. The IgJ^CreERT2^ model allowed for detailed analysis of surface marker expression on PCs, revealing insights into PC heterogeneity and characteristics. Our findings not only validate the IgJ^CreERT2^ mouse as a reliable tool for studying PCs but also facilitate the investigation of PC dynamics and longevity, particularly in the context of humoral immunity and vaccine responses. This model represents a significant advancement for the in-depth study of PCs in health and disease, offering a new avenue for the exploration of PC biology and immunological memory.

## Introduction

Plasma cells (PCs), highly specialized B cells, are critical to the humoral immune response. Following antigen exposure, activated B cells or reactivated memory B cells in germinal centers (GCs) or extrafollicular spaces can differentiate into PCs (Nutt, Hodgkin et al. 2015, Cyster and Allen 2019). This process is an essential part of the immune system that permits synthesis of immunoglobulins that specifically target pathogens and continues well after pathogen clearance. The environment influences PC function with factors such as cytokines and Toll-like receptor (TLR) ligands, leading to an increase in antibody production (Pioli 2019). PCs can exhibit other effector functions, including the production of cytokines such as interleukin(IL)-10, IL-17 and IL-35 which regulate immune responses to infections or progression to autoimmunity (Bermejo, Jackson et al. 2013, Shen, Roch et al. 2014, Rojas, Probstel et al. 2019). The multifaceted role of PCs makes them pivotal players in the immune landscape.

PCs serve as reservoirs of immunity, generating antigen-specific antibodies that offer protection against future exposures to pathogens (Nutt, Hodgkin et al. 2015, Schuh, Mielenz et al. 2020). This process, known as immunological memory, is the underlying basis of vaccination (Cyster and Allen 2019). The differentiation of B cells into PCs involves significant changes, including increased cytoplasmic volume and a rise in the number of endoplasmic reticulum and mitochondria; this facilitates the metabolic shift required to sustain elevated antibody secretion (Duan, Nguyen et al. 2023). This differentiation process relies on the coordination of several transcription factors that promote the expression of plasma cell-related genes while suppressing genes critical for maintaining B lymphocyte identity (Nutt, Hodgkin et al. 2015). To establish a PC-specific program, B lymphocyte-induced maturation protein (BLIMP1) functions as a transcriptional repressor of the B cell lineage transcription factor paired box protein 5 (PAX5) (Lin, Angelin-Duclos et al. 2002, Mikkola, Heavey et al. 2002, Shapiro-Shelef, Lin et al. 2003, Tellier, Shi et al. 2016). The repression of PAX5 then allows for the upregulation of X-box-binding protein (XBP1) and Interferon-regulatory factor 4 (IRF4) (Shaffer, Shapiro-Shelef et al. 2004, Low, Brodie et al. 2019). XBP1 regulates the unfolding protein response; this is crucial in preparing cells for large-scale antibody synthesis (Shaffer, Shapiro-Shelef et al. 2004). *IRF4* expression is vital for the differentiation and survival of PCs by supporting the PC transcriptional network, mitochondrial hemostasis and CD138 expression (Sciammas, Shaffer et al. 2006, Ochiai, Maienschein-Cline et al. 2013, Low, Brodie et al. 2019). This coordinated transcriptional reprogramming is necessary for the generation of antibody-secreting cells.

Upon exposure to their cognate antigen, B cells differentiate into PCs that undergo a series of epigenetic and transcriptional changes enabling them to survive for long periods (Brynjolfsson, Persson Berg et al. 2018, Cyster and Allen 2019). These cells can migrate to protective niches, such as bone marrow, and upregulate anti-apoptotic molecules while downregulating pro-apoptotic signals (Brynjolfsson, Persson Berg et al. 2018, Nguyen, Garimalla et al. 2018, Benet, Jing et al. 2021, Joyner, Ley et al. 2022). The distinction between short-lived plasma cells (SLPCs) and long-lived plasma cells (LLPCs) is particularly relevant to vaccination, allergy, immunological memory and immunity from natural infections. LLPCs, which can persist for months or even years within the bone marrow, differ from SLPCs by their enhanced survival capabilities (Brynjolfsson, Persson Berg et al. 2018, Benet, Jing et al. 2021). Investigating the generation and maintenance of LLPCs is a topic of great interest, however, it remains challenging due to the lack of distinguishing markers that differentiate LLPCs from SLPCs.

To advance our understanding of the fundamental biology of PCs, a lineage-specific Cre is essential for precise genetic studies. While transcription factors like BLIMP1, XPBP1, and IRF4 mediate terminal differentiation of B lymphocytes into PCs, they are not ideal for a plasma cell-specific mouse model due to their expression in other cell types (Kallies, Hawkins et al. 2006, Martins, Cimmino et al. 2006, Martinon, Chen et al. 2010, Man, Gabriel et al. 2017). Commonly used tools such as Blimp-1 reporters and Cre drivers have limitations, as Blimp-1 expression in a fraction of lymphocytes and myeloid cells introduces some limitations for this model (Xu, Barbosa et al. 2020, Nadeau and Martins 2022). In contrast, J-chain, a polypeptide essential for IgM and IgA oligomerization and normally repressed by PAX5 appears to be restricted to all PCs in mice irrespective of the immunoglobins they express (Rinkenberger, Wallin et al. 1996). This demonstrates promising characteristics for a lineage-specific Cre (Castro and Flajnik 2014). Given the importance of PCs in immunity, both our team and other researchers (Ayala, Bonaud et al. 2020, Xu, Barbosa et al. 2020), have generated a conditional J- chain EGPF CreERT2 mouse model. While the GFP protein is expressed in all J-chain expressing cells, upon tamoxifen administration, Cre-ERT2 is activated enabling time-specific genetic modifications (Indra, Warot et al. 1999).

To establish the utility of the J-chain CreERT2 GFP allele (IgJ^CreERT2^) mouse model for PC studies, we conducted a series of *in vitro* and *in vivo* validation studies utilizing the locus-encoded eGFP and crossed these mice with the tdTomato Cre-reporter strain to facilitate temporal tracking. PC-specific Cre recombinase expression is consistent with previously reported findings (Ayala, Bonaud et al. 2020, Xu, Barbosa et al. 2020). Using this model, we meticulously tracked PC responses, scrutinized the compartment-specific expression of the Cre allele, and validated the expression of commonly utilized PC surface markers. The temporal component and specificity afforded by this Cre recombinase makes this model particularly well-suited for the study of LLPCs in traditionally challenging tissue sites, such as the bone marrow and mucosal sites. Our studies highlight the utility of the IgJ^CreERT2^ mouse model as an invaluable tool for in-depth PC characterization in diverse physiological contexts, providing new insights into their behavior in the context of health and disease.

## Results

### Assessing the functionality of the J-chain Cre^ERT2^ ^GFP^ allele through *in vitro* studies

We derived a novel plasma cell-specific J-chain Cre (IgJCre^ERT2^), which was crossed to a tdTomato Cre reporter mouse to enable temporal labeling of J-Chain expressing PCs *in vitro* and *in vivo*. The Cre^ERT2^-p2A-GFP transgene is strategically targeted into the J-chain locus via an exon trap approach with an upstream splice acceptor site. The insertion ensures concurrent Cre and GFP expression whenever the J-chain allele is transcribed **Figure 1A**. Upon administration of tamoxifen, the Cre^ERT2^ protein undergoes nuclear translocation, inducing recombination and excision of a floxed stop cassette positioned upstream of the tdTomato allele. This removal of the stop cassette enables the subsequent expression of the tdTomato fluorescent protein.

**Figure 1.**
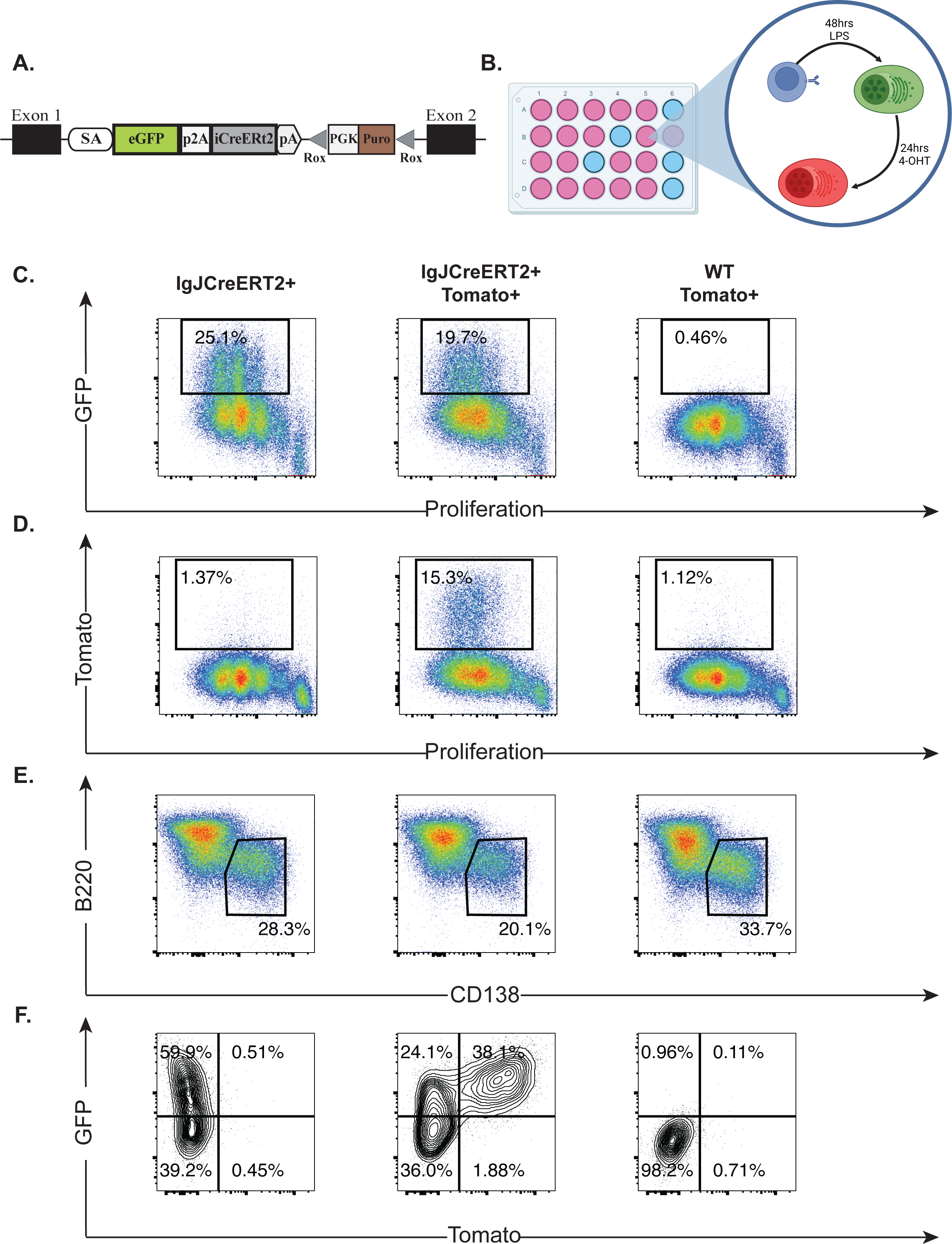
**A.** Schematic of the IgJCre^ERT2^ locus. **B.** Experimental setup for the *in vitro* differentiation of plasma cells using LPS and 4-Hydroxy-tamoxifen (4-OHT). B cells from IgJCre^ERT2^, IgJCre^ERT2^ tdTomato+, and littermate controls were isolated by CD43 depletion, stimulated with LPS, and evaluated by flow cytometry. Representative flow plots show: **C.** GFP fluorescence versus cell trace violet proliferation dye; **D.** tdTomato fluorescence versus cell trace violet proliferation dye. **E.** PCs were defined by CD138 expression and were further analyzed for **F.** GFP and tdTomato expression.

First, our initial objective was to validate the IgJCre^ERT2^ mouse model and that GFP expression was restricted to PCs. To differentiate PCs *in vitro*, we isolated mature B cells from the splenocytes of mice heterozygous for the IgJCre^ERT2^ allele or IgJCre^ERT2^ tdTomato^+^, or littermate controls. Subsequently, these cells were exposed to LPS for seventy-two hours. After forty-eight hours in culture with LPS, the cells were treated with 1000nM hydroxytamoxifen (4- OHT) to induce Cre-mediated recombination of the floxed tdTomato allele in IgJCre^ERT2^ expressing cells, **Figure 1B**. Upon LPS stimulation, B lymphocytes upregulated J-chain, with detectable GFP fluorescence after three rounds of cell division in IgJCre^ERT2^ positive cells, **Figure 1C**. Consistent with the cells that also harbored the tdTomato reporter and were exposed to 4-OHT, there was distinct red fluorescence after three rounds of cell division, **Figure 1D**. After seventy-two hours, about 60% of cells expressing CD138, the most routinely used surface marker for PCs, were also found to be GFP^+^, **Figures 1E, 1F**. For the IgJCre^ERT2-^ littermate controls, we used CD138 surface expression to define the PC population. In the absence of LPS, B cells cultured with the pro-survival cytokine BAFF did not proliferate or show any detectable CD138, GFP or tdTomato expression. Furthermore, the absence of tdTomato expression in conditions lacking 4-OHT in IgJCre^ERT2^ and CAG-tdTomato positive cells, illustrates the tight regulation of this novel tamoxifen-inducible Cre allele, **Supplementary Figure 1A**. The IgJCre^ERT2-^, tdTomato^+^cells did not have any GFP or tdTomato fluorescence even with 4-OHT exposure, **Figures 1C, 1D, 1F**. We observed a slight reduction in the CD138^+^ population for mice with the CreERT2 allele (p-value = 0.1061) suggesting slight Cre-associated toxicity, **Supplemental Figure 1B**. This toxicity was not observed in our *in vivo* experiments that we will discuss for the remainder of the manuscript, but underscores the importance of using appropriate Cre^ERT2-^ and CreERT2 controls while using this model. These results indicate the specificity of this labeling to both J-chain and CreERT2 expressing cells, allowing for successful cell-specific and temporal labeling of PCs *in vitro*.

### *In vivo* validation of the IgJ-Cre^ERT2^ allele following immunization

To test the application of the IgJCreERT2 mouse model *in vivo,* we employed sheep red blood cell (SRBC) immunizations, a T cell-dependent antigen known for inducing a robust GC response. The kinetics of the GC and PC responses to SRBC immunization were first tracked over a 60-day period in wild-type C57/BL6 mice in both the spleen and bone marrow using flow cytometry (**Supplementary Figure 2A**). PCs were identified by CD138^+^ expression and were either B220 low or high FSC in the spleen and bone marrow respectively. GC B cells were defined as CD38^-^ and FAS^+^, as depicted in the gating schemes in **Supplementary Figure 3**. In these experiments, we did not distinguish between plasmablast and PC populations as we did not stain for proliferation markers such as Ki67. We observed a strong GC response in the spleen that peaked at Day 10 post-initial SRBC immunization, **Supplementary Figure 2B**. There was an increase in splenic CD138^+^ cells between days 5-10, which declined thereafter, **Supplementary Figure 2C**. In the bone marrow, we did not observe any immunization-specific responses in CD138^+^ cells, **Supplementary Figure 2D**. These results emphasize how temporal labeling of PC populations would enhance our understanding of PC dynamics given the short-lived spleen and undetectable bone marrow PC responses. Labeling of the PC populations during the initial phase of the SRBC immunizations would permit us to assess their long-term maintenance and survival.

To address how effective the IgJCre^ERT2^ allele would be for lineage tracing of PCs, we repeated the SRBC immunizations in IgJCre^ERT2^ mice with and without the tdTomato reporter and compared them to CreERT2^-^ littermate controls. We immunized mice with SRBCs, and exposed mice to tamoxifen for five days between days 1 and 5, **Figure 2A**. Subsequently, mice were euthanized on day 5 and day 60 to evaluate the frequency of GFP^+^ and GFP^+^ tdTomato^+^ cells in their CD138^+^ population. We selected Day 5 as we aimed to monitor the initial stages of the PC response, building on our observations from **Supplemental Figure 2C**; and by day 60 we felt confident classifying any remaining GFP^+^Tomato^+^ cells as longer-lived PCs as they would have differentiated any time before or during tamoxifen administration.

**Figure 2.**
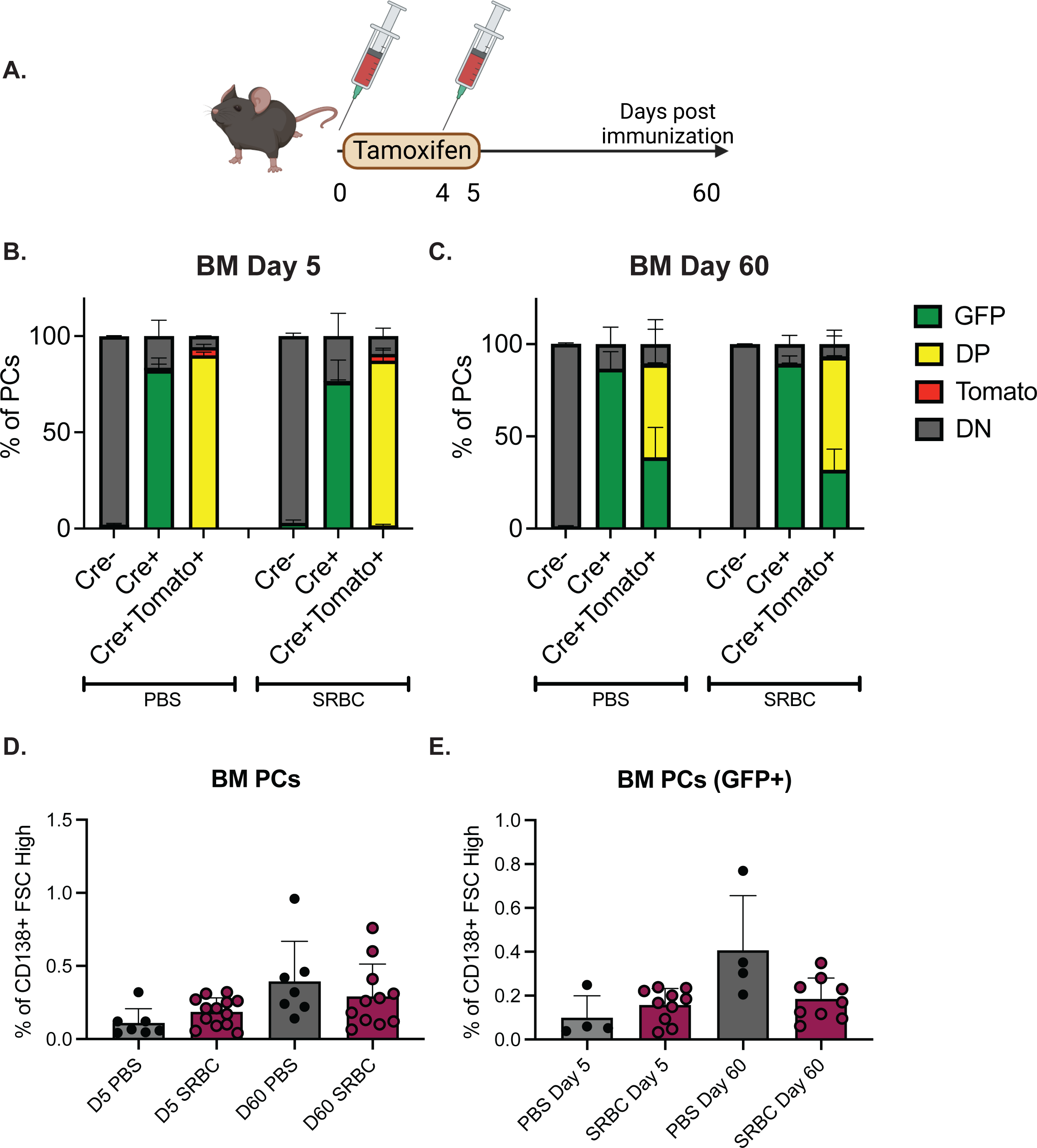
**A.** Schematic of SRBC and tamoxifen administration in mice. Percentage of bone marrow PCs at **B.** Day 5 and **C.** Day 60 defined as FSC^high^ CD138^+^ live singlets that were either GFP^+^, tdTomato^+^, GFP^+^tdTomato^+^ double positive (DP), or GFP^-^ tdTomato^-^ double negative (DN). The graphs summarize two independent experiments with 4-5 mice per group. **D.** Percentage of FSC^high^ CD138^+^ live singlets at days 5 and 60, separated by immunization status. **E.** Percentage of GFP^+^ FSC^high^ CD138^+^ live singlets at days 5 and 60, separated by immunization status.

Following tamoxifen administration, we observed 90% tdTomato positivity after gating on CD138^+^ and B220 low events in the spleen, and CD138^+^ and FSC-high events in the bone marrow (gating seen in **Supplementary Figure 3**). This was observed in both PBS and SRBC- immunized IgJCre^ERT2^ tdTomato^+^ mice at day 5 at both tissue sites, **Figure 2B, Supplementary Figure 4B**. At day 60 after tamoxifen labeling, only about 50-60% of PCs were still double-positive (DP) for GFP^+^ and tdTomato^+^ in both the PBS and SRBC-treated mice in both bone marrow, **Figure 2C**, and spleen **Supplementary Figure 4C**. SRBC immunization did not result in a significant increase in PCs within the bone marrow at either the early or late time point **Figure 2D**. This observation held when GFP expression alone was used to define the PC population in IgJ^CreERT2^ mice versus relying on the traditional CD138 surface expression **Figure 2E**. In the spleen, there was a clear increase at day 5 in both CD138^+^ and GFP^+^ PCs after SRBC immunization that was no longer present at day 60 **Supplementary Figures 4D, 4E**. At day 5, we observed a minor increase in GC B cells in the spleen of SRBC immunized mice **Supplementary Figure 4F**. Previous studies have shown low levels of *J-chain* transcripts in GC B cells, at least 40x less than PCs and J-chain protein expression in B220 high, CD138- GC B cells (Xu, Barbosa et al. 2020). In our SRBC immunization experiments, on day 5, we noted that approximately 15-20% of GC B cells displayed GFP expression, **Supplementary Figure 4G**, though this percentage did not change with immunization status of at day 60. Overall, we found that tamoxifen administration allows specific labeling of PCs in the spleen and bone marrow of IgJCre^ERT2^ tdTomato^+^ mice, and we did not observe any tdTomato expression when the mice only had the IgJCre^ERT2^ allele, **Figure 2** and **Supplementary Figure 4**. The fact that we did not observe a change in the frequency of PCs underscores the importance of labeling and tracking PCs that were present prior to/during immunization so that we can assess their longevity afterward.

### Plasma cell surface marker validation

CD138 is traditionally used as a surface marker to identify PCs, but its utility is hampered by several limitations. These include its susceptibility to collagenase cleavage (Goodyear, Kumar et al. 2014), expression on other cell types such as epithelial cells and fibroblasts, and CD138 shedding in malignant cells in multiple myeloma (Manon_JJensen, Itoh et al. 2010, Jung, Trapp-Stamborski et al. 2016). We observed that collagenase digestion of both small intestine (**Supplemental Figure 5A**) and lung tissue (**Supplemental Figure 5B**) revealed a GFP^+^ population with heterogeneity in CD138 staining. This variability could be indicative of non-specific CD138 cleavage. Additionally, the use of the IgJ^CreERT2^ GFP reporter was helpful for the lung tissue where PCs are in low abundance, and the GFP fluorescence enhanced the ability to identify this population by flow cytometry (**Supplemental Figure 5B**). For these reasons, the IgJCre^ERT2^ reporter model offers more confident identification of PCs. Using this novel model, we focused on examining the most utilized PC markers in GFP^+^ cells, following immunization with SRBCs. Our aim was to better understand the phenotypic heterogeneity in PCs in both spleen and bone marrow (Liu, Yao et al. 2022, Duan, Nguyen et al. 2023). This strategy simultaneously validated the IgJ^CreERT2^ model and confirmed the fidelity of some of the surface markers used to identify PCs in the absence of a reporter gene.

On day 5 following SRBC immunization, we isolated cells from the spleen and bone marrow of IgJCre^ERT2^ mice. Spleen was chosen to capture the SLPC response, and bone marrow as the site where we anticipated more LLPCs and biological heterogeneity in surface marker expression. After gating on CD45^+^ FSC-High GFP^+^ live-singlets, we analyzed the expression of CD19, CD138, MHCII, CD44, CD98, CD69, and CD93 on PCs and compared them to B220^+^ B cells. In both spleen and bone marrow, PCs, defined by GFP expression, were positive for CD44, CD98 and CD138, as depicted in **Figure 3A and 3B**. Conversely, PCs at both sites displayed a downregulation of CD19, and CD93 expression varied between splenic PCs (bimodal) and bone marrow PCs (spectrum of expression), as depicted in **Figure 3**. Splenic PCs exhibited positivity for MHCII but were negative for CD69 expression (**Figure 3A**). In contrast, bone marrow PCs showed higher CD69 expression compared to B2 cells or splenic PCs and displayed variable MHCII expression (**Figure 3B**). Our findings not only corroborate the presence of well-established markers like CD138 and CD98 in GFP^+^ cells but also offer a more nuanced assessment of the expression of other surface markers, including CD19, CD44, CD69, CD93 and MHCII. Together, these results enrich our understanding of the phenotypic characteristics of GFP^+^ PC, shedding light on the heterogeneity of surface marker expression, and underscore the utility of the IgJCre^ERT2^ reporter mouse.

**Figure 3.**
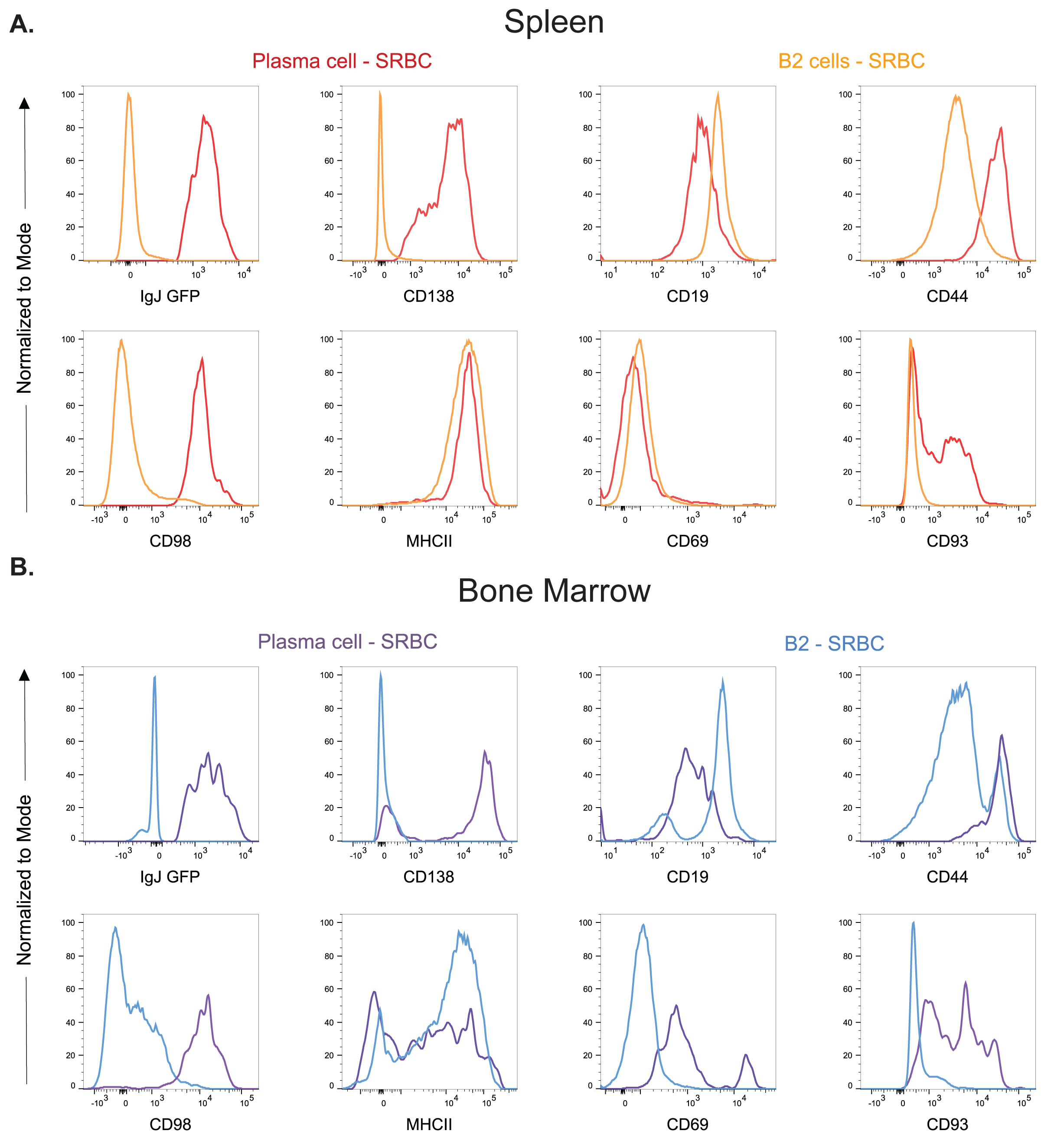
IgJCre^ERT2^ mice were immunized with SRBCs. On day 5, the **A.** spleen and **B**. bone marrow were collected and GFP^+^ live CD45^+^ singlets were evaluated for their expression of GFP, CD138, CD19, CD44, CD98, MHCII, CD69 and CD93 and compared to B2 cells, defined as CD19/B220^+^ cells.

## Discussion

PCs play a pivotal role in the humoral immune response due to their ability to establish enduring antibody-mediated immunity, produce cytokines, and respond to TLR signaling. Nevertheless, the limited tools to study these cells *in vivo* has impeded research on these versatile cells. To overcome this challenge, we have established a novel inducible PC-specific Cre driver, the IgJ^CreERT2^ mouse. This innovative tool enables temporal tracking and genetic editing of PCs, unlocking new avenues for comprehensive investigations into the intricate PC biology.

The IgJ model offers numerous advantages for tracking PCs. For instance, the reliance on CD138, a common PC marker, is known to be sensitive to trypsin cleavage (Liu and Akkoyunlu 2021) and collagenase cleavage (Schaffer, Maul-Pavicic et al. 2019) complicating its use in tissue digestion protocols, **Supplemental Figure 5**. The loss of CD138 expression in the presence of sodium azide (Wilmore, Jones et al. 2017), and rapid loss of CD138 expression *ex vivo* within 30 minutes after isolation further complicates its use (Dang, Mohr et al. 2022). We have found the IgJ^CreERT2^ mouse emerges as a solution to overcome the limitations, providing a stable and reliable alternative with GFP as a tracking marker evident in **Supplemental Figure 5**. GFP proves to be a more stable and reliable marker for tracking PCs, emphasizing its utility, esecially in tissues that require enzymatic digestion. Thus, the model holds significant potential for advancing the characterization of PCs in disease, exploring homeostatic diversity, and addressing challenges related to technical sample collection.

We utilized this mouse model to track the GC and PC response and turnover *in vivo* following immunization with SRBCs, a commonly utilized T-cell dependent immunization protocol. While SRBC immunizations are frequently used, the actual data detailing the kinetics of the PC responses is sparse, with more emphasis on GC responses in existing literature. Our GC findings are consistent with previous reports on GC dynamics. For instance, studies by McAllister, Apgar et al. (2017) identified SRBC-specific splenic GC B cells at day 7, and Zhang, Tech et al. (2018) used IHC to monitor the splenic GC B cells over time, and found them as early as day 4 through day 14. Our data, **Supplemental Figures 2B, 4F, 4G**, aligns with these observations, where we saw GCs in the spleen increase at our initial time point of day 5 and peak at day 10. Our experimental design allowed us to monitor PC kinetics *in vivo* during SRBC immunization, shedding light on long-term PC responses. McAllister, Apgar et al. (2017) found SRBC-specific antibodies in serum as early as day 7, the earlier time point of their study, suggesting that PCs were already producing SRBC-specific antibodies by this time. Others assessed PC responses by IF staining for IRF4, and found PCs could be observed as early as day 3, peaked at days 5-6, and dropped again by day 9 (Zhang, Tech et al. 2018). This is consistent with our findings following SRBC immunizations, **Supplemental Figure 2C**. Notably, SRBC immunization did not significantly change the percentage of PCs in the bone marrow at either day 5 or 60 (**Figure 2D and 2E**).

The turnover of PCs in the spleen and bone marrow was assessed at day 60 by the frequency of GFP^+^ and tdTomato^+^ PCs. Any PCs formed after tamoxifen labeling will be GFP^+^ tdTomato^-^, while any PCs that were 60 days or older would be tdTomato^+^. We observed comparable frequencies of tdTomato^+^ PCs in the bone marrow (∼60%), regardless of whether or not the animals were immunized with SRBC (**Figure 2C**). In the spleen at day 60, we noted that the frequency of tdTomato+ cells had decreased from 90% to ∼25%, indicating a higher rate of PC turnover in the spleen than in the bone marrow (**Supplemental Figure 4B and 4C**). Others reported a bone marrow PC half-life of ∼200 days, and observed similar spleen PC kinetics to what we found in our experiments (Xu, Barbosa et al. 2020). However, the kinetics of PCs responses to other antigens, vaccines, and in the steady state have not been thoroughly investigated. This gap in knowledge is partly attributable to T cell-dependent PCs exhibiting low levels of membrane-bound BCRs, which hinders tracking of antigen-specific PCs (Blanc, Moro-Sibilot et al. 2016). The IgJ^CreERT2^ model offers a potential solution to some of the technical challenges associated with tracking PC responses, facilitating the study of PC kinetics, along with genetic manipulation of these cells.

The IgJ^CreERT2^ model allowed us to undertake the characterization of PC surface markers in both spleen and bone marrow during SRBC immunization. Our examination encompassed the previously established PC surface markers, including CD138, CD98, and CD44 in both the spleen and bone marrow (**Figure 3**), aligning with prior studies and corroborating existing descriptions by others (Cassese, Arce et al. 2003, Tellier and Nutt 2017, Dang, Mohr et al. 2022). While other research groups have reported heterogeneity in PCs across various tissues (Wilmore, Gaudette et al. 2021, Joyner, Ley et al. 2022, Liu, Yao et al. 2022), we aimed to explore additional PC surface marker expression, specifically CD93, MHCII, and CD69, in the spleen and bone marrow (**Figure 3**). In our study, the majority of splenic PCs did not express CD69, whereas in the bone marrow, PC exhibited variable CD69 expression. This variability points to the potential utility of CD69 as a distinguishing marker for bone marrow-resident PCs, similar to its established role in lung-resident memory B cells (Barker, Etesami et al. 2021). Given that recent PC activation in the bone marrow is unlikely, CD69 expression may contribute to the maintenance of PC longevity in the bone marrow niche. For MHCII expression, we observed it predominately in B cells and PCs in the spleen, which likely represents newly minted PCs or PBs. Conversely, the bone marrow PCs showed a spectrum of MHCII expression, the MHCII low cells being indicative of a more mature PC population (Manz, Thiel et al. 1997, Slifka, Antia et al. 1998). The investigation into CD93 revealed a mixed population in both spleen and bone marrow, with a slightly higher proportion in the bone marrow. This aligns with previous research which linked CD93 to the maintenance of antibody secretion and PC retention in the bone marrow (Chevrier, Genton et al. 2009). Together, these findings underscore the heterogeneity of PCs. Our model offers a new avenue for characterizing PC surface markers and their lifespan across different tissues, enhancing our understanding of the nuanced roles that these cells play beyond antibody secretion.

In conclusion, the IgJ^CreERT2^ mouse model is a genetic tool that facilitates the study of PCs with greater precision and detail. This model overcomes previous technical barriers, enabling tracking and genetic profiling of PCs. Our work not only validates this model, but also expands the knowledge on PC biology, including response kinetics, turnover, and surface marker characterization during SRBC immunization. This model’s potential to elucidate the diverse functions and subpopulations of PCs promises to foster new insights into their role in immunity and disease.

### Limitations of the Study

Early Cre^ERT2^ and GFP expression in the GC cells prior to production of transcriptional (BCL6^-^, IRF4^+^, Blimp1^+^) and phenotypic markers of PCs (CD138^+^, FAS^-^, CD38^+^) may limit the applications of the IgJ^CreERT2^ model for the genetic study of malignant transformation of PCs (MGUS and multiple myeloma) as the Cre expression will occur earlier in the developmental lineage of PCs than would be desirable. Also, although the model provides invaluable genetic “timestamping”, its applicability for different tissues and across different immunizations remains to be fully optimized and explored. Furthermore, tamoxifen induction of Cre^ERT2^ can have off target physiological effects that need to be controlled for appropriately in all studies. Finally, IgJ expression in humans is restricted to IgA and IgM-producing PCs and this is not mirrored in mice. This species-specific different in the regulation of J-chain allele enables broader application of the Cre model, but users should be aware of this difference. Overall, the utility of this Cre driver paves the way for numerous genetic investigations into PC biology.

## Acknowledgements

The authors thank Eric Bartnicki for technical input and Dr. Susan Schwab’s lab for advice regarding tamoxifen administration.

## Author Contributions

Conceptualization, TCB, KZ, AMZ, MJH, SBK. Formal analysis, TCB, KZ, AMZ, SBK. Investigation, TCB, KZ, AMZ, MJH, SB, TM, TB, TH. Writing-original draft, TCB, KZ, AMZ, KC, SBK. Writing-review and editing, TCB, KZ, AMZ, MJH, VH, KC, SBK; Supervision, KC, SBK. Funding acquisition, KC, SBK.

## Materials and Methods

### Mice

C57BL/6J and tdTomato (strain #:007914) mice were obtained from Jackson Laboratories and housed at New York University School of Medicine. The Wellcome Trust Sanger Institute generated the J-chain CreERT2 allele. C57BL/6 J-Chain CreERT2 mouse embryonic stem cells were purchased from the European Mouse Mutant Archive (EMMA) and rederived at NYU School of Medicine. The J-Chain CreERT2 and tdTomato lines were maintained on a C57BL/6J background. All mouse experiments followed federal and institutional regulations, approved by the New York University Langone Institutional Animal Care and Use Committee (IACUC protocol IA16-01399). Mice had ad libitum access to food and were maintained on a 12-hour light-dark schedule.

### Flow Cytometry

Spleen and bone marrow were harvested from mice at euthanasia. The spleen was mechanically disrupted over a 70uM filter using the back of syringe in RPMI (Corning: 10- 040-CV) supplemented with 10% fetal bovine serum and 1X penicillin and streptomycin (complete RPMI). The bone marrow was flushed out of a decapped femur using a 27G needle, and 3mL of complete RPMI, and immediately resuspended using 1 mL pipette to generate a single cell suspension. Both spleen and bone marrow were subjected to red blood cell lysis using Pharm Lyse (BD: 555899). Single-cell suspensions were then stained with fluorescently tagged antibodies for evaluation on a BD Fortessa. Data was analyzed using FlowJo Version 10.9 (BD)

### SRBC Immunizations and Tamoxifen Administration

Single-cell suspensions of spleens were prepared and subjected to CD43 depletion (Invitrogen: 11422D). Mature B cells at >90% purity were stained with cell trace violet (Invitrogen: C34557) and subjected to plasma cell differentiation using 10ug/mL of LPS (Sigma: L4391-1MG) for 72 hours. After 48 hours in culture with LPS, 1000nM of (Z)-4-Hydroxytamoxifen was added to the culture (Sigma: H7904- 5MG). Cells were analyzed using flow cytometry on a BD Fortessa.

### SRBC Immunizations and Tamoxifen Administration

Sterile-defibrinated SRBCs were purchased from Cedar Lane (CL2581-100D). For immunizations, SRBCs were washed twice by topping up the volume to 50mL with sterile phosphate buffered solution (PBS, Corning: 21-040- CV) and spun down for 10 minutes at 3000RPM. SRBCs were then counted using a hemocytometer, and the cell density was adjusted to 5×10^9^ cells/mL. Then 1×10^9^ SRB cells were injected into the mouse on days 0 and 4. 100mg of Tamoxifen (Sigma: T5648-1G) was brought to room temperature and subjected to end over end mixing in 5mL of corn oil at 37C until dissolved. Then 100uL of tamoxifen was administered to mice each day for five consecutive days (2mg per day).

### Statistical Analyses

We performed statistical analysis using Graphpad Prism 10.0.1 software. To test for statistical significance, we used a Mann-Whitney test or Kruskal-Wallis test (one-way non-parametric ANOVA). We considered differences statistically significant when p < 0.05.

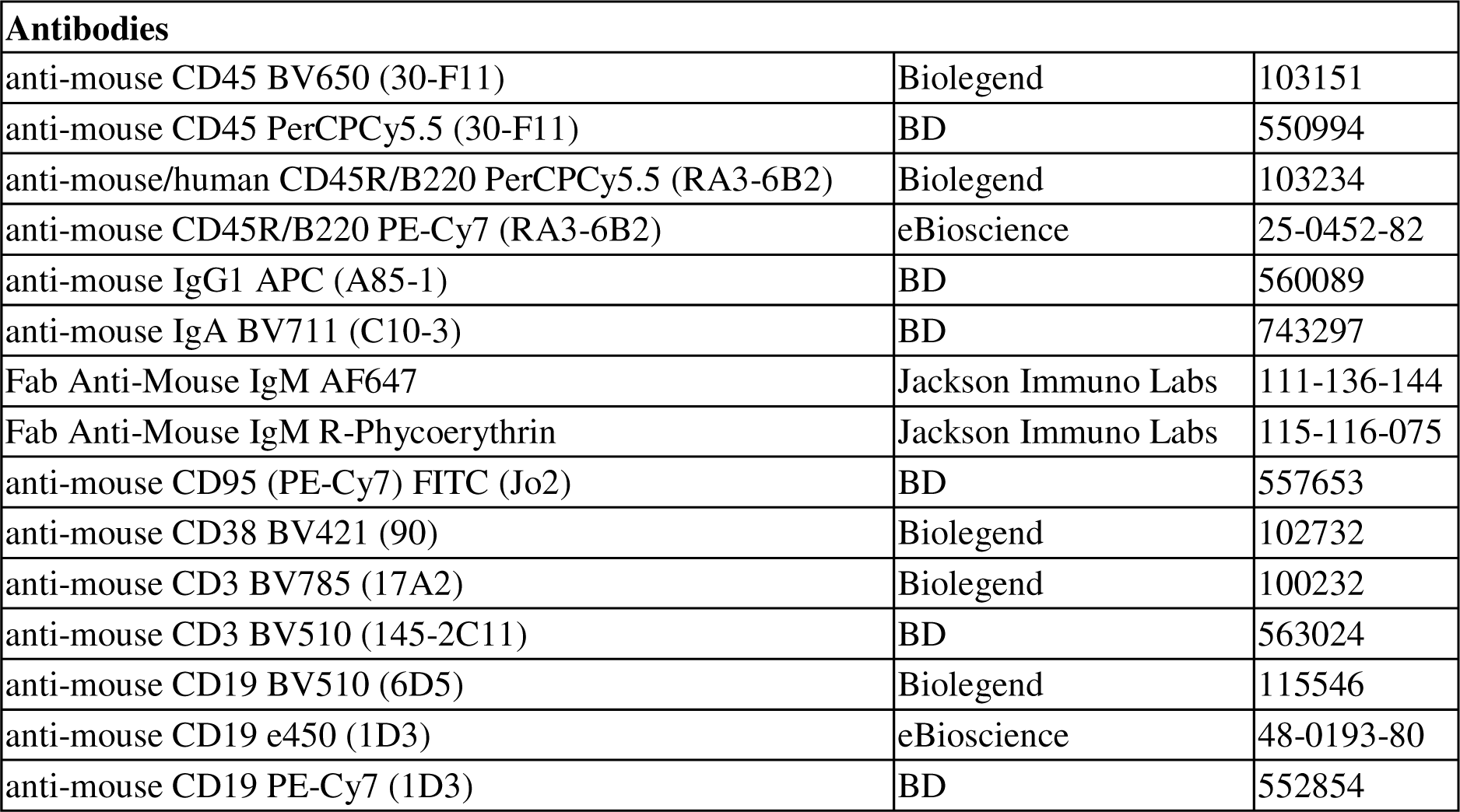
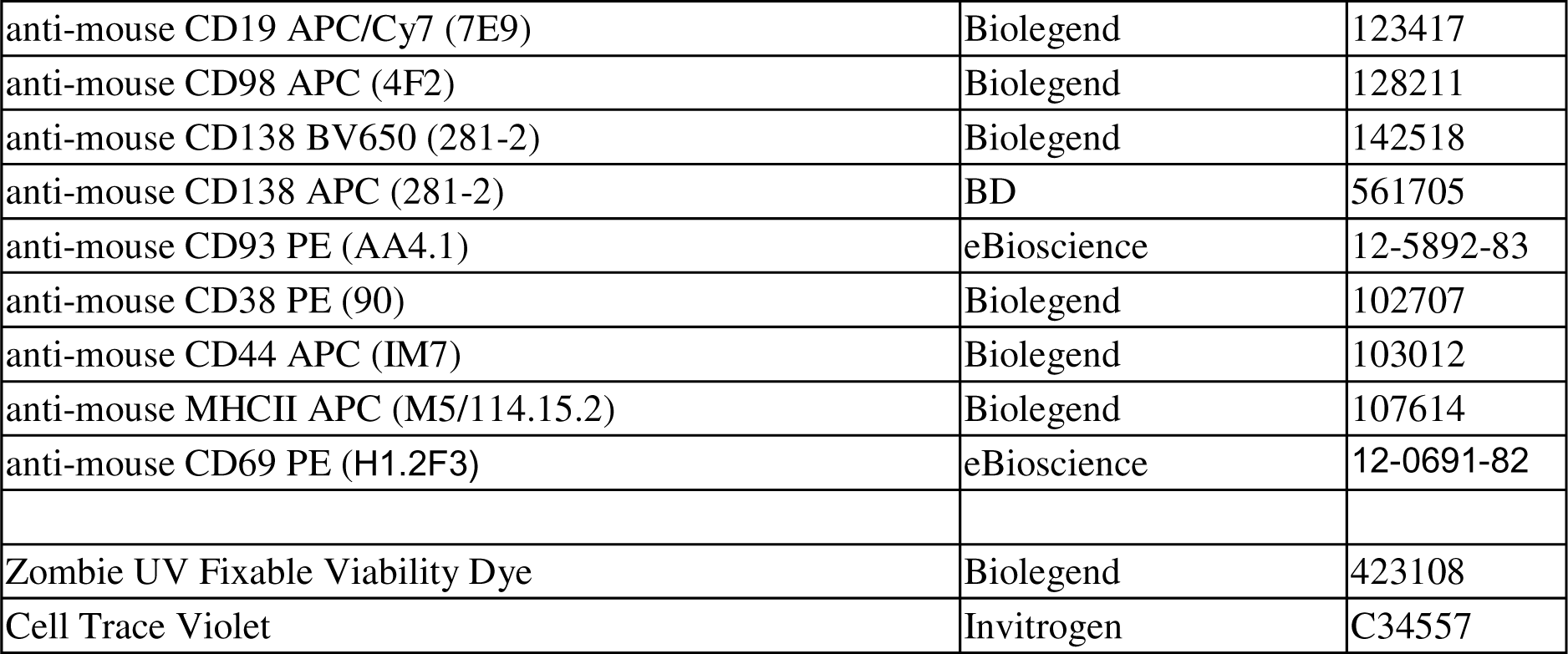

---

**Supplemental Figure 1.**
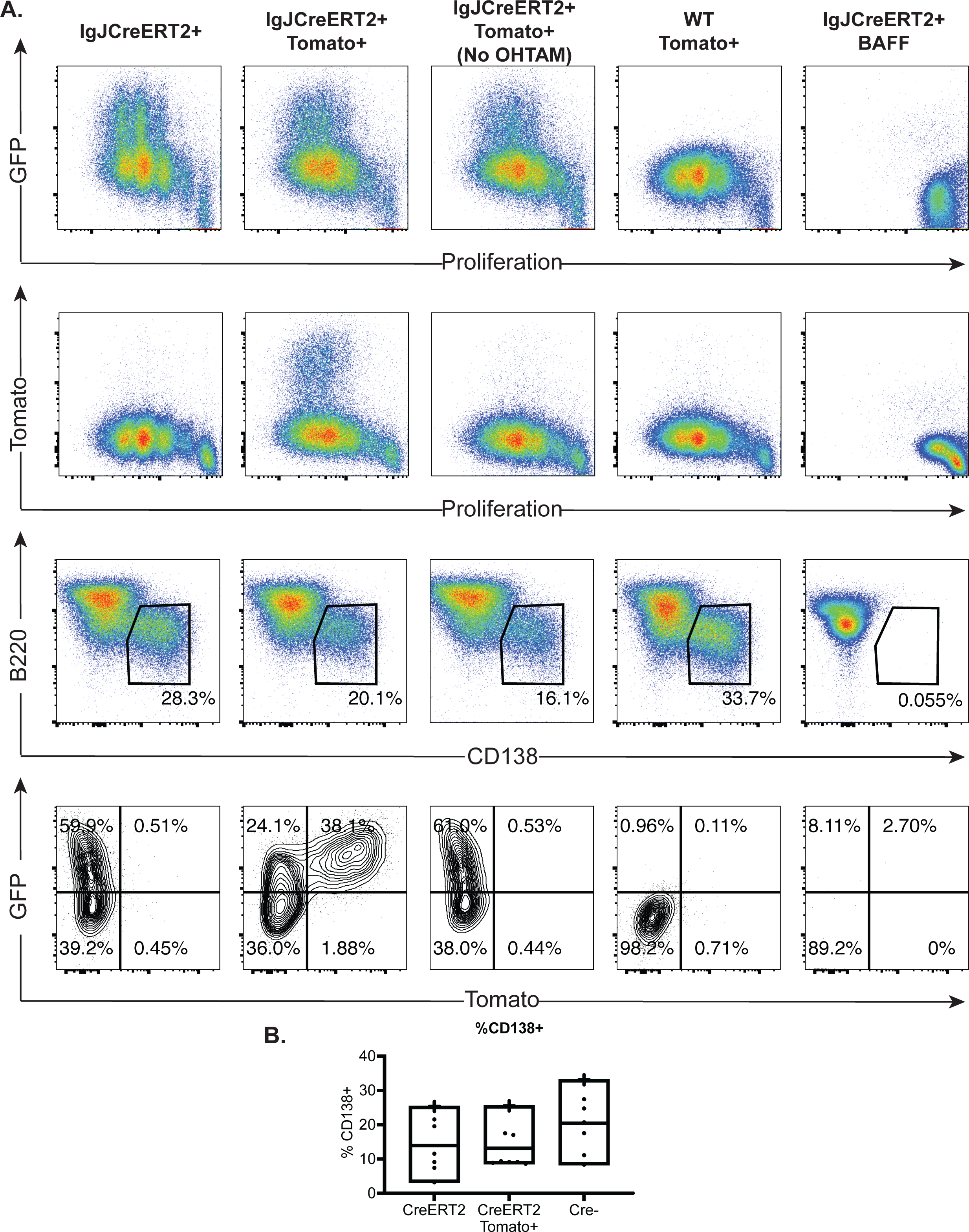
**A.** B cells from IgJCre^ERT2^, IgJCre^ERT2^ tdTomato^+^, and littermate controls were isolated via CD43 depletion and stimulated with LPS, or BAFF. Following flow cytometry analysis, cells were evaluated for GFP fluorescence versus Cell Trace Violet proliferation dye, tdTomato fluorescence versus Cell Trace Violet proliferation dye, and plasma cells were identified by CD138 expression and further analyzed for GFP and Tomato expression. **B.** Data from multiple *in vitro* experiments in which CD43 depleted B cells from IgJCre^ERT2^, IgJCre^ERT2^ tdTomato^+^, and littermate controls were stimulated with LPS. Each data point is the average of 2-3 technical replicates.

**Supplemental Figure 2.**
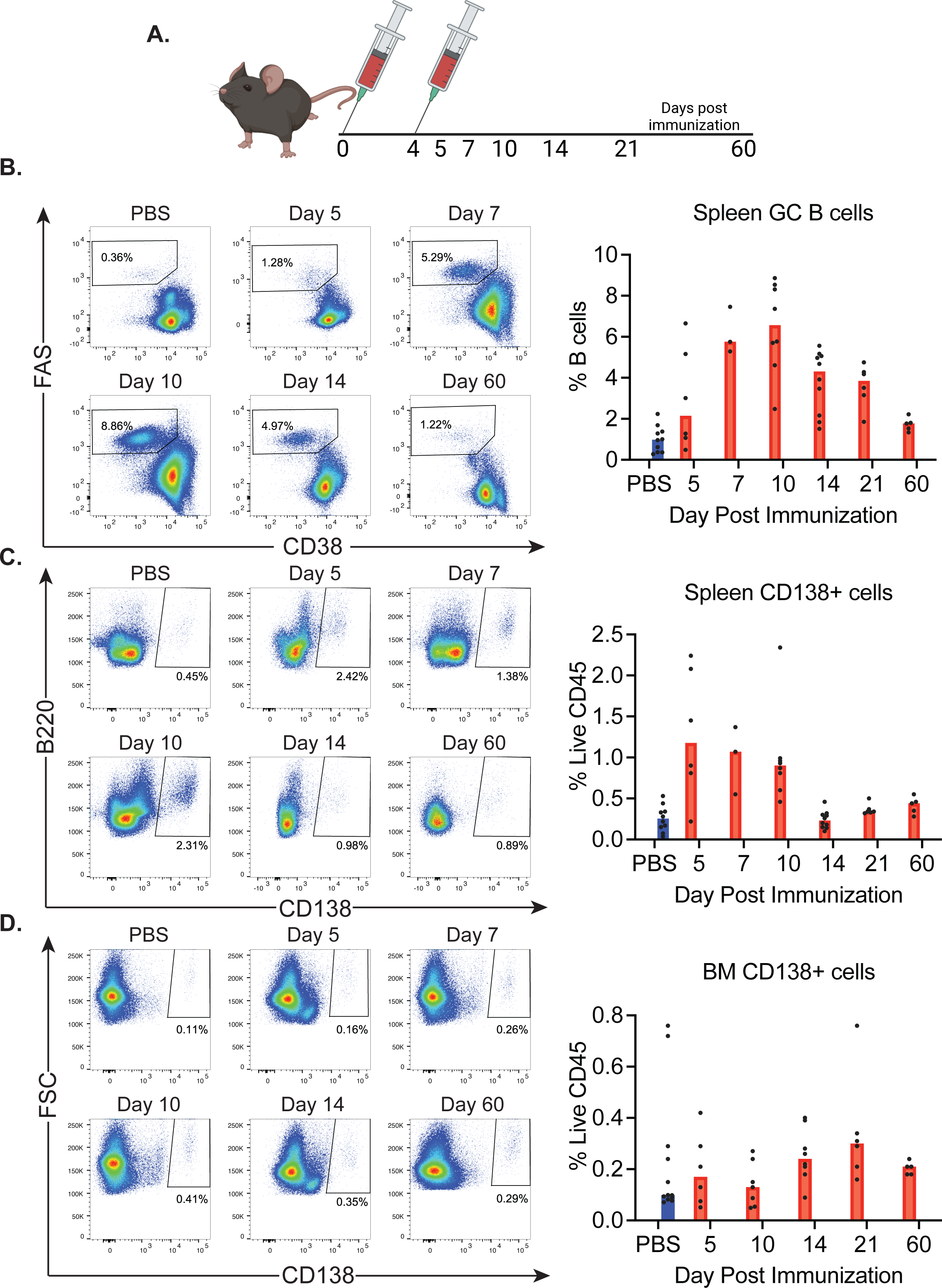
**A.** C57BL/6J mice were immunized with SRBCs and spleen and bone marrow were evaluated by flow cytometry on days 5, 7, 10, 14, 21 and 60. **B.** Representative flow cytometry plots and the summary for spleen GC B cell frequencies. **C.** Representative flow cytometry plots and summary for splenic CD138^+^ PCs. **D.** Representative flow cytometry plots of bone marrow CD138^+^ PCs.

**Supplemental Figure 3.**
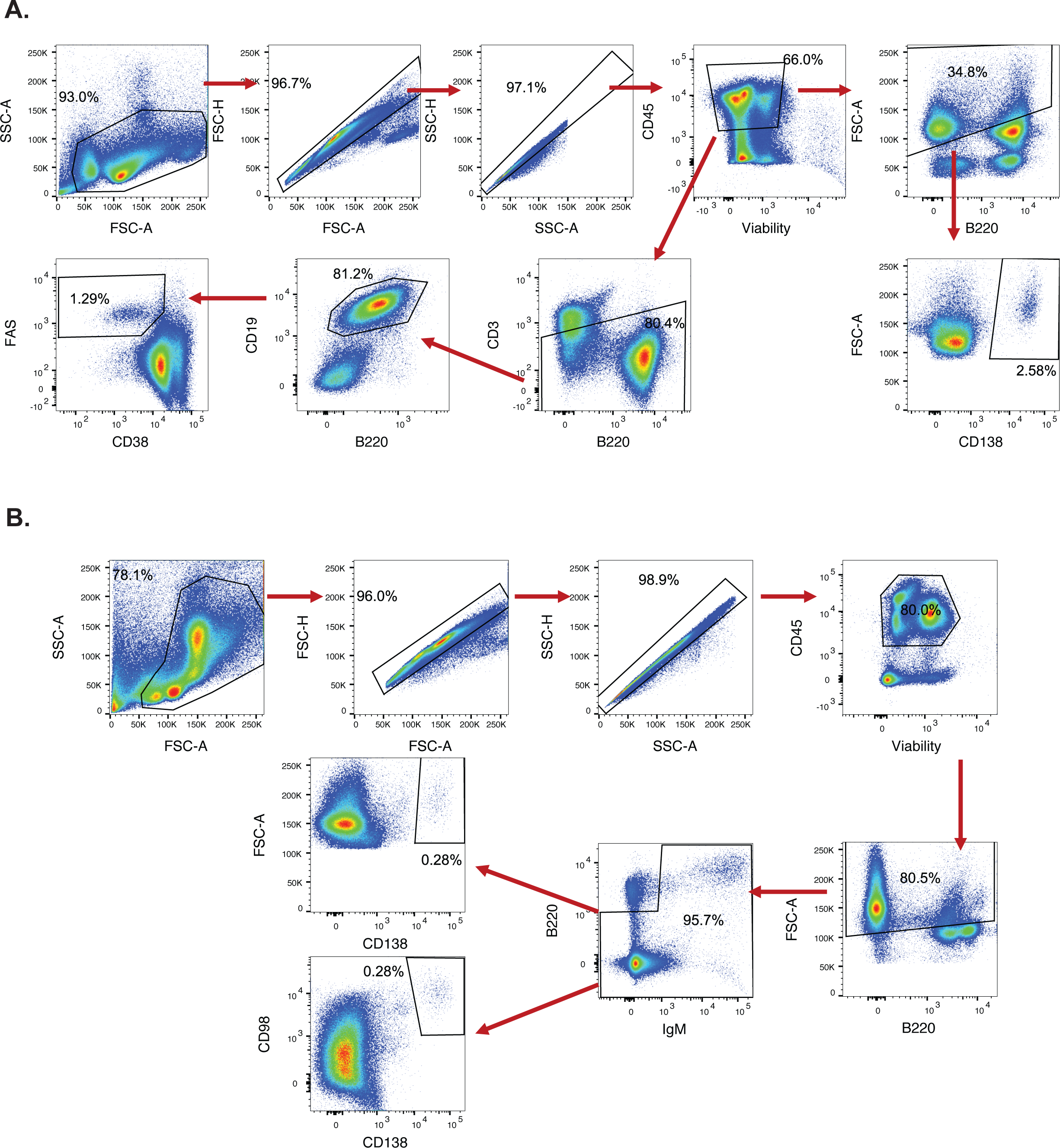
**A.** Gating strategy for the analysis of spleen by flow cytometry. **B.** Gating strategy for the analysis of bone marrow by flow cytometry.

**Supplemental Figure 4.**
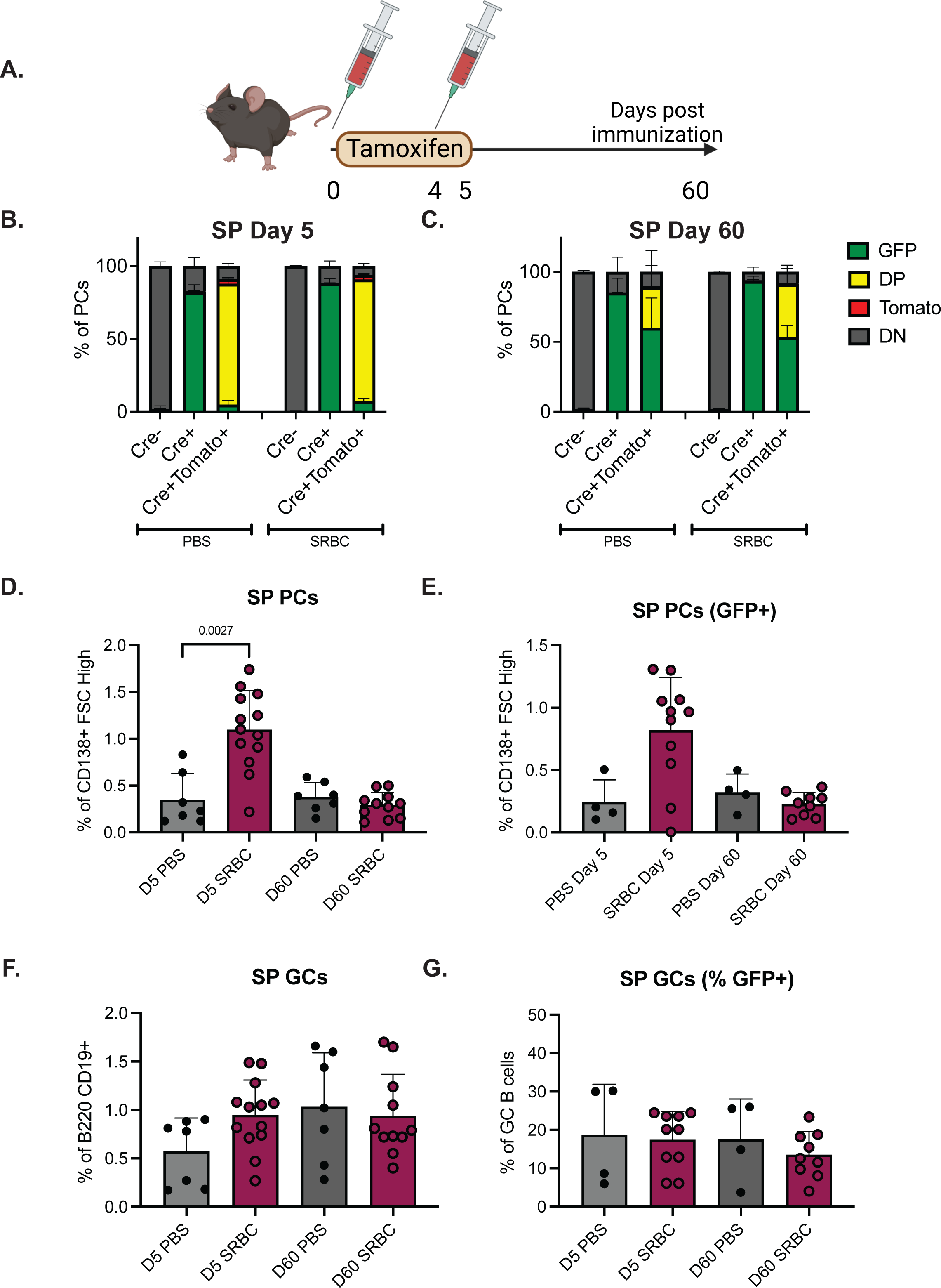
**A.** Schematic of SRBC and tamoxifen administration in mice. Percent of splenic plasma cells at **B.** Day 5 and **C.** Day 60 defined as FSC^high^ CD138^+^ live singlets that are GFP^+^, tdTomato^+^, GFP^+^tdTomato^+^ double positive (DP), or GFP^-^ tdTomato^-^ double negative (DN). Graphs are the summary of two independent experiments with 4-5 mice per group. **D.** The percentage of FSC^high^ CD138^+^ live singlets at days 5 and 60, separated by immunization status. **E.** The percentage of GFP^+^ FSC^high^ CD138^+^ live singlets at days 5 and 60, separated by immunization status. **F.** Spleen GCs at day 5 and 60 represented as percent of B2 cells. **G.** The percentage of GC B cells that are GFP^+^ at day 5 and 60 separated by immunization status.

**Supplemental Figure 5.**
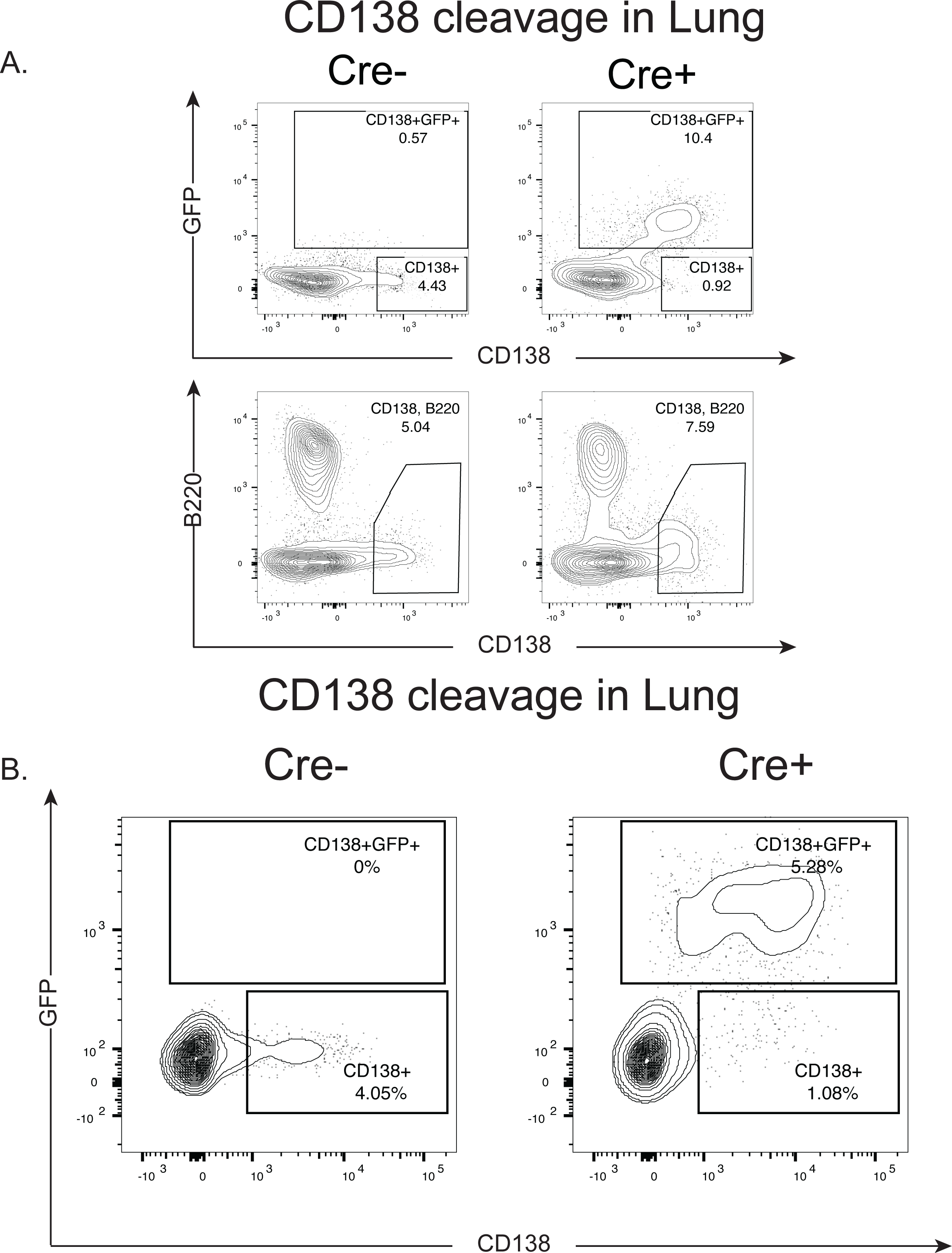
**A.** IgJCre^ERT2^ and littermate control small intestinal tissue was digested using collagenase and DNaseI to isolate cells from the lamina propria. These cells were evaluated by flow cytometry. **B.** D IgJCre^ERT2^ and littermate control lung tissue was digested using collagenase and DNaseI to isolate cells and these cells were evaluated by flow cytometry.

## References

Ayala, M. V., A. Bonaud, S. Bender, J.-M. Lambert, F. Lechouane, C. Carrion, M. Cogné, V. Pascal and C. Sirac (2020). “New models to study plasma cells in mouse based on the restriction of IgJ expression to antibody secreting cells.” bioRxiv: 2020.2008.2013.249441.

Barker, K. A., N. S. Etesami, A. T. Shenoy, E. I. Arafa, C. Lyon de Ana, N. M. Smith, I. M. Martin, W. N. Goltry, A. M. Barron, J. L. Browning, H. Kathuria, A. C. Belkina, A. Guillon, X. Zhong, N. A. Crossland, M. R. Jones, L. J. Quinton and J. P. Mizgerd (2021). “Lung-resident memory B cells protect against bacterial pneumonia.” J Clin Invest 131(11).

Benet, Z., Z. Jing and D. R. Fooksman (2021). “Plasma cell dynamics in the bone marrow niche.” Cell Reports 34(6): 108733.

Bermejo, D. A., S. W. Jackson, M. Gorosito-Serran, E. V. Acosta-Rodriguez, M. C. Amezcua-Vesely, B. D. Sather, A. K. Singh, S. Khim, J. Mucci, D. Liggitt, O. Campetella, M. Oukka, A. Gruppi and D. J. Rawlings (2013). “Trypanosoma cruzi trans-sialidase initiates a program independent of the transcription factors RORgammat and Ahr that leads to IL-17 production by activated B cells.” Nat Immunol 14(5): 514–522.

Blanc, P., L. Moro-Sibilot, L. Barthly, F. Jagot, S. This, S. de Bernard, L. Buffat, S. Dussurgey, R. Colisson, E. Hobeika, T. Fest, M. Taillardet, O. Thaunat, A. Sicard, P. Mondière, L. Genestier, S. L. Nutt and T. Defrance (2016). “Mature IgM-expressing plasma cells sense antigen and develop competence for cytokine production upon antigenic challenge.” Nat Commun 7: 13600.

Brynjolfsson, S. F., L. Persson Berg, T. Olsen Ekerhult, I. Rimkute, M.-J. Wick, I.-L. Mårtensson and O. Grimsholm (2018). “Long-Lived Plasma Cells in Mice and Men.” Frontiers in Immunology 9.

Cassese, G., S. Arce, A. E. Hauser, K. Lehnert, B. Moewes, M. Mostarac, G. Muehlinghaus, M. Szyska, A. Radbruch and R. A. Manz (2003). “Plasma cell survival is mediated by synergistic effects of cytokines and adhesion-dependent signals.” J Immunol 171(4): 1684–1690.

Castro, C. D. and M. F. Flajnik (2014). “Putting J chain back on the map: how might its expression define plasma cell development?” J Immunol 193(7): 3248–3255.

Chevrier, S., C. Genton, A. Kallies, A. Karnowski, L. A. Otten, B. Malissen, M. Malissen, M. Botto, L. M. Corcoran, S. L. Nutt and H. Acha-Orbea (2009). “CD93 is required for maintenance of antibody secretion and persistence of plasma cells in the bone marrow niche.” Proc Natl Acad Sci U S A 106(10): 3895–3900.

Cyster, J. G. and C. D. C. Allen (2019). “B Cell Responses: Cell Interaction Dynamics and Decisions.” Cell 177(3): 524–540.

Dang, V. D., E. Mohr, F. Szelinski, T. A. Le, J. Ritter, T. Hinnenthal, A. L. Stefanski, E. Schrezenmeier, S. Ocvirk, C. Hipfl, S. Hardt, Q. Cheng, F. Hiepe, M. Lohning, T. Dorner and A. C. Lino (2022). “CD39 and CD326 Are Bona Fide Markers of Murine and Human Plasma Cells and Identify a Bone Marrow Specific Plasma Cell Subpopulation in Lupus.” Front Immunol 13: 873217.

Duan, M., D. C. Nguyen, C. J. Joyner, C. L. Saney, C. M. Tipton, J. Andrews, S. Lonial, C. Kim, I. Hentenaar, A. Kosters, E. Ghosn, A. Jackson, S. Knechtle, S. Maruthamuthu, S. Chandran, T. Martin, R. Rajalingam, F. Vincenti, C. Breeden, I. Sanz, G. Gibson and F. E. Lee (2023). “Understanding heterogeneity of human bone marrow plasma cell maturation and survival pathways by single-cell analyses.” Cell Rep 42(7): 112682.

Goodyear, A. W., A. Kumar, S. Dow and E. P. Ryan (2014). “Optimization of murine small intestine leukocyte isolation for global immune phenotype analysis.” Journal of Immunological Methods 405: 97–108.

Indra, A. K., X. Warot, J. Brocard, J.-M. Bornert, J.-H. Xiao, P. Chambon and D. Metzger (1999). “Temporally-controlled site-specific mutagenesis in the basal layer of the epidermis: comparison of the recombinase activity of the tamoxifen-inducible Cre-ERT and Cre-ERT2 recombinases.” Nucleic Acids Research 27(22): 4324–4327.

Joyner, C. J., A. M. Ley, D. C. Nguyen, M. Ali, A. Corrado, C. Tipton, C. D. Scharer, T. Mi, M. C. Woodruff, J. Hom, J. M. Boss, M. Duan, G. Gibson, D. Roberts, J. Andrews, S. Lonial, I. Sanz and F. E.-H. Lee (2022). “Generation of human long-lived plasma cells by developmentally regulated epigenetic imprinting.” Life Science Alliance 5(3): e202101285.

Jung, O., V. Trapp-Stamborski, A. Purushothaman, H. Jin, H. Wang, R. Sanderson and A. Rapraeger (2016). “Heparanase-induced shedding of syndecan-1/CD138 in myeloma and endothelial cells activates VEGFR2 and an invasive phenotype: prevention by novel synstatins.” Oncogenesis 5(2): e202–e202.

Kallies, A., E. D. Hawkins, G. T. Belz, D. Metcalf, M. Hommel, L. M. Corcoran, P. D. Hodgkin and S. L. Nutt (2006). “Transcriptional repressor Blimp-1 is essential for T cell homeostasis and self-tolerance.” Nat Immunol 7(5): 466–474.

Lin, K.-I., C. Angelin-Duclos, T. C. Kuo and K. Calame (2002). “Blimp-1-dependent repression of Pax-5 is required for differentiation of B cells to immunoglobulin M-secreting plasma cells.” Molecular and cellular biology 22(13): 4771–4780.

Liu, L. and M. Akkoyunlu (2021). “Circulating CD138 enhances disease progression by augmenting autoreactive antibody production in a mouse model of systemic lupus erythematosus.” J Biol Chem 297(3): 101053.

Liu, X., J. Yao, Y. Zhao, J. Wang and H. Qi (2022). “Heterogeneous plasma cells and long-lived subsets in response to immunization, autoantigen and microbiota.” Nature Immunology 23(11): 1564–1576.

Low, M. S. Y., E. J. Brodie, P. L. Fedele, Y. Liao, G. Grigoriadis, A. Strasser, A. Kallies, S. N. Willis, J. Tellier, W. Shi, S. Gabriel, K. O’Donnell, C. Pitt, S. L. Nutt and D. Tarlinton (2019). “IRF4 Activity Is Required in Established Plasma Cells to Regulate Gene Transcription and Mitochondrial Homeostasis.” Cell Rep 29(9): 2634–2645.e2635.

Man, K., S. S. Gabriel, Y. Liao, R. Gloury, S. Preston, D. C. Henstridge, M. Pellegrini, D. Zehn, F. Berberich-Siebelt, M. A. Febbraio, W. Shi and A. Kallies (2017). “Transcription Factor IRF4 Promotes CD8(+) T Cell Exhaustion and Limits the Development of Memory-like T Cells during Chronic Infection.” Immunity 47(6): 1129–1141.e1125.

Manon_JJensen, T., Y. Itoh and J. R. Couchman (2010). “Proteoglycans in health and disease: the multiple roles of syndecan shedding.” The FEBS journal 277(19): 3876–3889.

Manz, R. A., A. Thiel and A. Radbruch (1997). “Lifetime of plasma cells in the bone marrow.” Nature 388(6638): 133–134.

Martinon, F., X. Chen, A. H. Lee and L. H. Glimcher (2010). “TLR activation of the transcription factor XBP1 regulates innate immune responses in macrophages.” Nat Immunol 11(5): 411–418.

Martins, G. A., L. Cimmino, M. Shapiro-Shelef, M. Szabolcs, A. Herron, E. Magnusdottir and K. Calame (2006). “Transcriptional repressor Blimp-1 regulates T cell homeostasis and function.” Nat Immunol 7(5): 457–465.

McAllister, E. J., J. R. Apgar, C. R. Leung, R. C. Rickert and J. Jellusova (2017). “New Methods To Analyze B Cell Immune Responses to Thymus-Dependent Antigen Sheep Red Blood Cells.” J Immunol 199(8): 2998–3003.

Mikkola, I., B. Heavey, M. Horcher and M. Busslinger (2002). “Reversion of B cell commitment upon loss of Pax5 expression.” Science 297(5578): 110–113.

Nadeau, S. and G. A. Martins (2022). “Conserved and Unique Functions of Blimp1 in Immune Cells.” Frontiers in Immunology 12.

Nguyen, D. C., S. Garimalla, H. Xiao, S. Kyu, I. Albizua, J. Galipeau, K.-Y. Chiang, E. K. Waller, R. Wu, G. Gibson, J. Roberson, F. E. Lund, T. D. Randall, I. Sanz and F. E.-H. Lee (2018). “Factors of the bone marrow microniche that support human plasma cell survival and immunoglobulin secretion.” Nature Communications 9(1): 3698.

Nutt, S. L., P. D. Hodgkin, D. M. Tarlinton and L. M. Corcoran (2015). “The generation of antibody-secreting plasma cells.” Nat Rev Immunol 15(3): 160–171.

Ochiai, K., M. Maienschein-Cline, G. Simonetti, J. Chen, R. Rosenthal, R. Brink, A. S. Chong, U. Klein, A. R. Dinner and H. Singh (2013). “Transcriptional regulation of germinal center B and plasma cell fates by dynamical control of IRF4.” Immunity 38(5): 918–929.

Pioli, P. D. (2019). “Plasma Cells, the Next Generation: Beyond Antibody Secretion.” Front Immunol 10: 2768.

Rinkenberger, J. L., J. J. Wallin, K. W. Johnson and M. E. Koshland (1996). “An interleukin-2 signal relieves BSAP (Pax5)-mediated repression of the immunoglobulin J chain gene.” Immunity 5(4): 377–386.

Rojas, O. L., A. K. Probstel, E. A. Porfilio, A. A. Wang, M. Charabati, T. Sun, D. S. W. Lee, G. Galicia, V. Ramaglia, L. A. Ward, L. Y. T. Leung, G. Najafi, K. Khaleghi, B. Garcillan, A. Li, R. Besla, I. Naouar, E. Y. Cao, P. Chiaranunt, K. Burrows, H. G. Robinson, J. R. Allanach, J. Yam, H. Luck, D. J. Campbell, D. Allman, D. G. Brooks, M. Tomura, R. Baumann, S. S. Zamvil, A. Bar-Or, M. S. Horwitz, D. A. Winer, A. Mortha, F. Mackay, A. Prat, L. C. Osborne, C. Robbins, S. E. Baranzini and J. L. Gommerman (2019). “Recirculating Intestinal IgA-Producing Cells Regulate Neuroinflammation via IL-10.” Cell 176(3): 610–624 e618.

Schaffer, S., A. Maul-Pavicic, R. E. Voll and N. Chevalier (2019). “Optimized isolation of renal plasma cells for flow cytometric analysis.” J Immunol Methods 474: 112628.

Schuh, W., D. Mielenz and H.-M. Jäck (2020). Chapter Three - Unraveling the mysteries of plasma cells. Advances in Immunology. F. W. Alt, Academic Press. 146: 57–107.

Sciammas, R., A. Shaffer, J. H. Schatz, H. Zhao, L. M. Staudt and H. Singh (2006). “Graded expression of interferon regulatory factor-4 coordinates isotype switching with plasma cell differentiation.” Immunity 25(2): 225–236.

Shaffer, A., M. Shapiro-Shelef, N. N. Iwakoshi, A.-H. Lee, S.-B. Qian, H. Zhao, X. Yu, L. Yang, B. K. Tan and A. Rosenwald (2004). “XBP1, downstream of Blimp-1, expands the secretory apparatus and other organelles, and increases protein synthesis in plasma cell differentiation.” Immunity 21(1): 81–93.

Shapiro-Shelef, M., K.-I. Lin, L. J. McHeyzer-Williams, J. Liao, M. G. McHeyzer-Williams and K. Calame (2003). “Blimp-1 is required for the formation of immunoglobulin secreting plasma cells and pre-plasma memory B cells.” Immunity 19(4): 607–620.

Shen, P., T. Roch, V. Lampropoulou, R. A. O’Connor, U. Stervbo, E. Hilgenberg, S. Ries, V. D. Dang, Y. Jaimes, C. Daridon, R. Li, L. Jouneau, P. Boudinot, S. Wilantri, I. Sakwa, Y. Miyazaki, M. D. Leech, R. C. McPherson, S. Wirtz, M. Neurath, K. Hoehlig, E. Meinl, A. Grutzkau, J. R. Grun, K. Horn, A. A. Kuhl, T. Dorner, A. Bar-Or, S. H. E. Kaufmann, S. M. Anderton and S. Fillatreau (2014). “IL-35-producing B cells are critical regulators of immunity during autoimmune and infectious diseases.” Nature 507(7492): 366–370.

Slifka, M. K., R. Antia, J. K. Whitmire and R. Ahmed (1998). “Humoral immunity due to long-lived plasma cells.” Immunity 8(3): 363–372.

Tellier, J. and S. L. Nutt (2017). “Standing out from the crowd: How to identify plasma cells.” Eur J Immunol 47(8): 1276–1279.

Tellier, J., W. Shi, M. Minnich, Y. Liao, S. Crawford, G. K. Smyth, A. Kallies, M. Busslinger and S. L. Nutt (2016). “Blimp-1 controls plasma cell function through the regulation of immunoglobulin secretion and the unfolded protein response.” Nature Immunology 17(3): 323–330.

Wilmore, J. R., B. T. Gaudette, D. Gómez Atria, R. L. Rosenthal, S. K. Reiser, W. Meng, A. M. Rosenfeld, E. T. Luning Prak and D. Allman (2021). “IgA Plasma Cells Are Long-Lived Residents of Gut and Bone Marrow That Express Isotype- and Tissue-Specific Gene Expression Patterns.” Front Immunol 12: 791095.

Wilmore, J. R., D. D. Jones and D. Allman (2017). “Protocol for improved resolution of plasma cell subpopulations by flow cytometry.” Eur J Immunol 47(8): 1386–1388.

Xu, A. Q., R. R. Barbosa and D. P. Calado (2020). “Genetic timestamping of plasma cells in vivo reveals tissue-specific homeostatic population turnover.” eLife 9: e59850.

Zhang, Y., L. Tech, L. A. George, A. Acs, R. E. Durrett, H. Hess, L. S. K. Walker, D. M. Tarlinton, A. L. Fletcher, A. E. Hauser and K.-M. Toellner (2018). “Plasma cell output from germinal centers is regulated by signals from Tfh and stromal cells.” Journal of Experimental Medicine 215(4): 1227–1243.

